# Chemical Induction of Longevity-Promoting Colanic Acid in the Host’s Microbiota

**DOI:** 10.1101/2024.07.23.604802

**Authors:** Guo Hu, Matthew Brandon Cooke, Alice X Wen, Xin Yu, Jin Wang, Christophe Herman, Meng C. Wang

## Abstract

Microbiota-derived metabolites have emerged as key regulators of longevity. The metabolic activity of the gut microbiota, influenced by dietary components and ingested chemical compounds, profoundly impacts host fitness. While the benefits of dietary prebiotics are well-known, chemically targeting the gut microbiota to enhance host fitness remains largely unexplored. Here, we report a novel chemical approach to induce a pro-longevity bacterial metabolite in the host gut. We discovered that specific *Escherichia coli* strains overproduce colanic acids (CAs) when exposed to a low dose of cephaloridine, leading to an increased lifespan in host *Caenorhabditis elegans*. In the mouse gut, oral administration of low-dose cephaloridine induces the transcription of the capsular biosynthesis operon responsible for CA biosynthesis in commensal *E. coli*, which overcomes the inhibition of CA biosynthesis above 30°C and enables its induction directly from the microbiota. Importantly, low-dose cephaloridine induces CA independently of its antibiotic properties through a previously unknown mechanism mediated by the membrane-bound histidine kinase ZraS. Our work lays the foundation for microbiota-based therapeutics through the chemical modulation of bacterial metabolism and reveals the promising potential of bacteria-targeting drugs in promoting host longevity.

## Main

### Introduction

The gut microbiota present in the gastrointestinal (GI) tract plays a crucial role in human health and disease susceptibility^1^, influencing the host’s neuronal functions^2^, immunity^3^, and life expectancy^4^. The microbiota genetic composition (termed microbiome) of over 2000 genera^5^ encodes crucial enzymes that mediate the production of unique microbial metabolites, host-derived secondary metabolites, and host-microbe-shared metabolites^6^. These microbial metabolites provide energy to local intestinal epithelial cells as essential nutrients^7^ and influence remote organs through blood circulation as signaling molecules^8^. The microbiota also responds to environmental inputs, such as changes in diet and medication uses, to produce bioactive compounds based on ingested diets^9^ or alter drug efficacy to be more or less active-even toxic^10^, resulting in both advantageous and disadvantageous outcomes for the host.

In recent years, an increasing number of microbiome-based therapeutics have shown benefits to the host, including fecal microbiota transplantation, dietary prebiotics, enteral reconstitution of symbiotic bacteria, the introduction of engineered bacteria, and supplementation of microbiota-derived bioactive compounds^11^. Here, we introduce a microbiome-based approach that uses host-impermeable drugs to specifically target metabolic biosynthesis pathways in gut commensals to promote host longevity. Distinct from those existing strategies, this approach chemically targets the existing commensal community to induce the production of metabolic products beneficial to the host. We demonstrated this approach in both *Caenorhabditis elegans* (*C. elegans*) and *Mus musculus* (mice), and further uncovered the molecular mechanism underlying this chemical-induced effect in *Escherichia coli* (*E. coli*).

## Results

### Chemically inducing colanic acid beyond temperature restriction to promote longevity

Colanic acid (CA) is an extracellular polysaccharide synthesized in *E. coli* and other *Enterobacteriaceae*^12^. Its biosynthesis is catalyzed by a series of enzymes encoded in the capsular biosynthesis (*cps*) operon^13^. Our previous studies have revealed the longevity-promoting effect of CA in *C. elegans* and *Drosophila melanogaster*^14,15^. Currently, its application in mammals is challenging because CA cannot be synthesized chemically or purified from *E. coli* in large quantities, nor can it be effectively delivered through the GI tract. It is known that *E. coli* is typically among the first commensal bacteria to inhabit the human gut^16^. However, CA production from *E. coli* becomes restricted when the environmental temperature exceeds 30°C^17^. A previous study indicated that certain β-lactam antibiotics trigger the overexpression of the *cps* operon in *E. coli* at sub-minimum inhibitory concentrations (sub-MIC) at 37°C^18^. We thus utilized an *E. coli* strain that harbors a LacZ reporter for the *cps* operon transcriptional regulation to screen multiple β-lactam antibiotics and antibiotics that disrupt the outer membrane, protein synthesis, and genome replication. Using colorimetric assay, we examined the levels of β-galactosidase from the transcriptional up-regulation of the *cps* operon, resulting in intensified blue coloration. We confirmed that cephaloridine (Cepha) and cefazolin (Cafez) strongly induce the *cps* operon, while cephalothin (CephaT), cefuroxime (Cefu), and penicillins-benzylpenicillin (PenG), ampicillin (Amp), and carbenicillin (Carb), and colistin show various levels of moderate effects based on the blue coloration (**Fig 1A** and **Supplementary Fig 1**).

**Fig 1.**
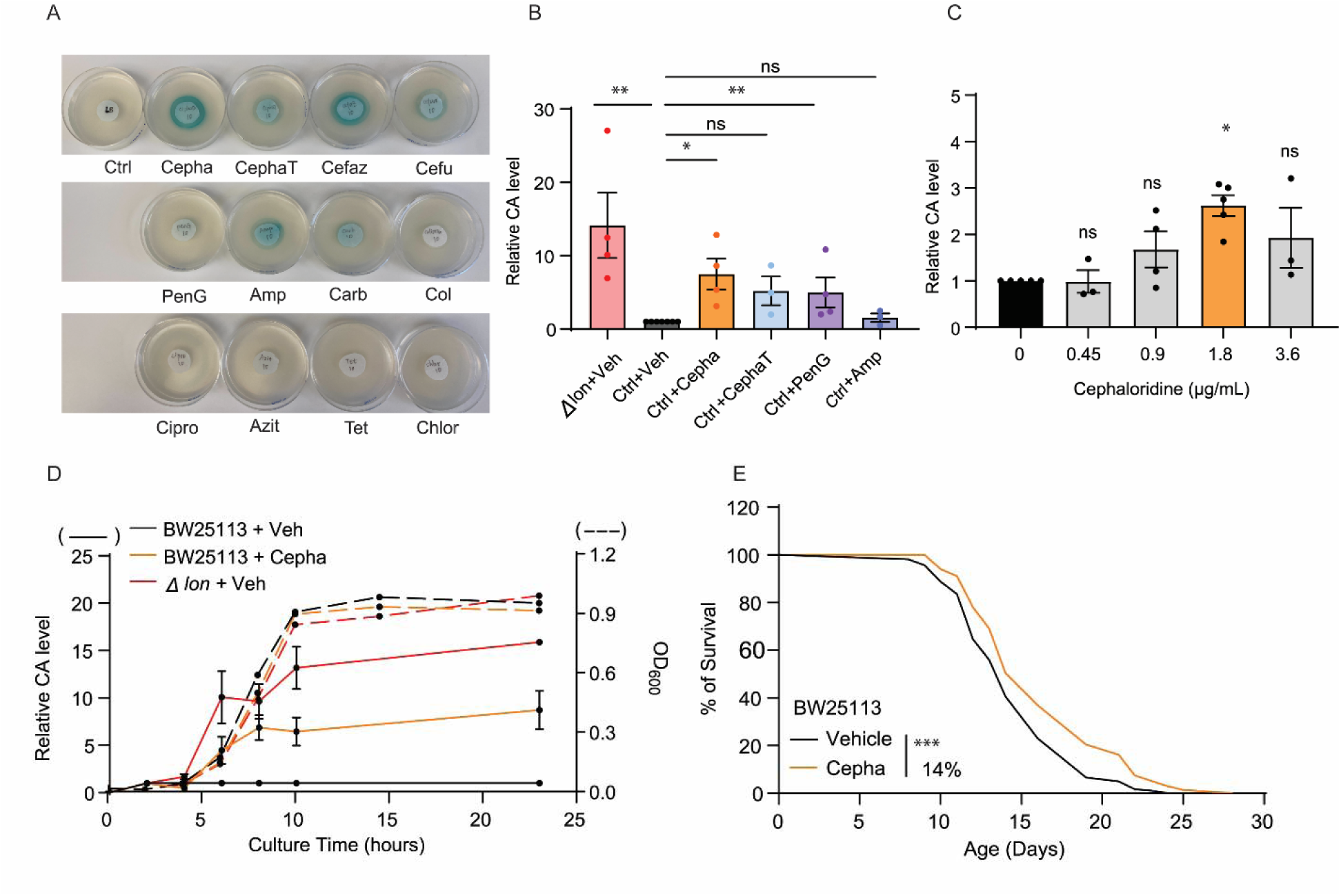
Chemical induction of bacterial CA promotes longevity in host *C. elegans*. (A) The disk-diffusion assay shows *E. coli* treated by chemical compounds presenting various blue colorations, indicating the induction of the *cps* operon responsible for colanic acid biosynthesis. Ctrl (LB medium), Cepha (cephaloridine), CephaT (cephalothin), Cefaz (Cefazolin), Cefu (cefuroxime), PenG (benzylpenicillin), Amp (ampicillin), Carb (carbenicillin), Col (colistin), Cipro (ciprofloxacin), Azit (Azithromycin), Tet (tetracycline), and Chlor (chloramphenicol); N=2. (B) Quantification of colanic acids (CA) from filtered bacterial cultures shows the *lon* deletion mutant *E. coli* (Δ*lon*) produces an increased level of CA by 14-fold as compared to the wild-type (WT) BW25113 *E. coli*. Cepha at 1.8 μg/mL and PenG at 5 μg/mL induce CA production by 7-fold and 5-fold, respectively; CephaT at 0.625 μg/mL and Amp at 1.5 μg/mL do not induce CA significantly (*p>0.05).* N ≥ 3 (C) Quantification of the relative CA induction in WT BW25113 *E. coli* when responding to 0, 0.45, 0.9, 1.8, and 3.6 μg/mL Cepha. A low dose of cephaloridine at 1.8 μg/mL is defined as the Colanic Acid-Inducing dose (Cepha-CAI dose), N ≥ 3 (D) Bacterial growth curves for WT BW25113 *E. coli* with the vehicle and Cepha-CAI dose treatments and the Δ*lon* mutant. WT *E. coli* treated by Cepha-CAI dose and the Δ*lon* mutant increase CA levels during the log phase (solid lines) and do not affect bacterial growth (dotted lines). N=2. (E) The lifespan of *C. elegans* is extended by 14% with Cepha-CAI dose of WT BW25113 *E. coli*. N = 3, 60 - 100 worms per replicate. In (B) and (C), error bars represent the mean ± standard error of the mean (s.e.m.). *\*\*p < 0.01, *p < 0.05, ns p > 0.05* by two-tailed Student’s t-test. In (E), *\*\*\*p < 0.001* by log-rank test.

Next, we quantified bacterial CA production in the bacteria culture medium treated with different antibiotics. The medium from *E. coli* treated with cephaloridine at sub-MIC showed the highest CA level, while penicillins and cephalothin showed lower or no increase (**Fig1B**). Furthermore, we found that the most effective CA-inducing concentration of cephaloridine was 1.8 μg/mL, which is lower than previously reported MIC values at either 2 μg/mL^19^ or 4 μg/mL^20^ in *E. coli* (**Fig 1C**). We referred to this low dose of cephaloridine as the Colanic Acid-Inducing dose (Cepha-CAI dose). Unlike the delayed growth, a common effect of sub-MIC antibiotic exposure^21^, wild-type *E. coli* treated with Cepha-CAI dose exhibited similar growth rates to those without the treatment and to the CA-overproducing Δ*lon* mutants^14^ (**Fig 1D**). Together, these results demonstrate that the Cepha-CAI dose overrides the temperature restriction of CA production at 37 °C without inhibiting *E. coli* growth or causing cell death.

We further revealed that similar to the Δ*lon* mutant, wild-type *E. coli* treated by Cepha-CAI dose showed increased CA production during the log phase, and the elevated levels persisted into the stationary phase (**Fig 1D**). Importantly, wild-type *C. elegans* with *E. coli* treated with the Cepha-CAI dose showed a 14% extension in lifespan compared to the controls without the treatment (**Fig 1E** and **Supplementary Table 1**). This demonstrates that chemically induced CA effectively confers longevity-promoting benefits in the host *C. elegans*.

### Low-dose cephaloridine induces CA biosynthesis in the mammalian gut

Due to its poor oral absorption, cephaloridine as an antibiotic agent was terminated in the clinic^25^. Even at a concentration of 1 mM, cephaloridine cannot be sufficiently absorbed by the GI tract due to a high efflux rate ^25^. Since we successfully induced CAs using the Cepha-CAI dose that bypassed the temperature constraint at 37°C, we hypothesized that cephaloridine would specifically target commensal *E. coli* and stimulate CA production in the mammalian gut, where the average body temperature ranges from 36°C to 40°C. To test this hypothesis, we first treated MG1655, a K-12 *E. coli* strain derived from human microbiota, with the Cepha-CAI dose at 37°C and detected increased CA levels in the culture medium (**Fig 2A**). Furthermore, wild-type *C. elegans* hosting the Cepha-CAI dose-treated MG1655 showed a 13% lifespan extension (**Fig 2B**).

**Fig 2.**
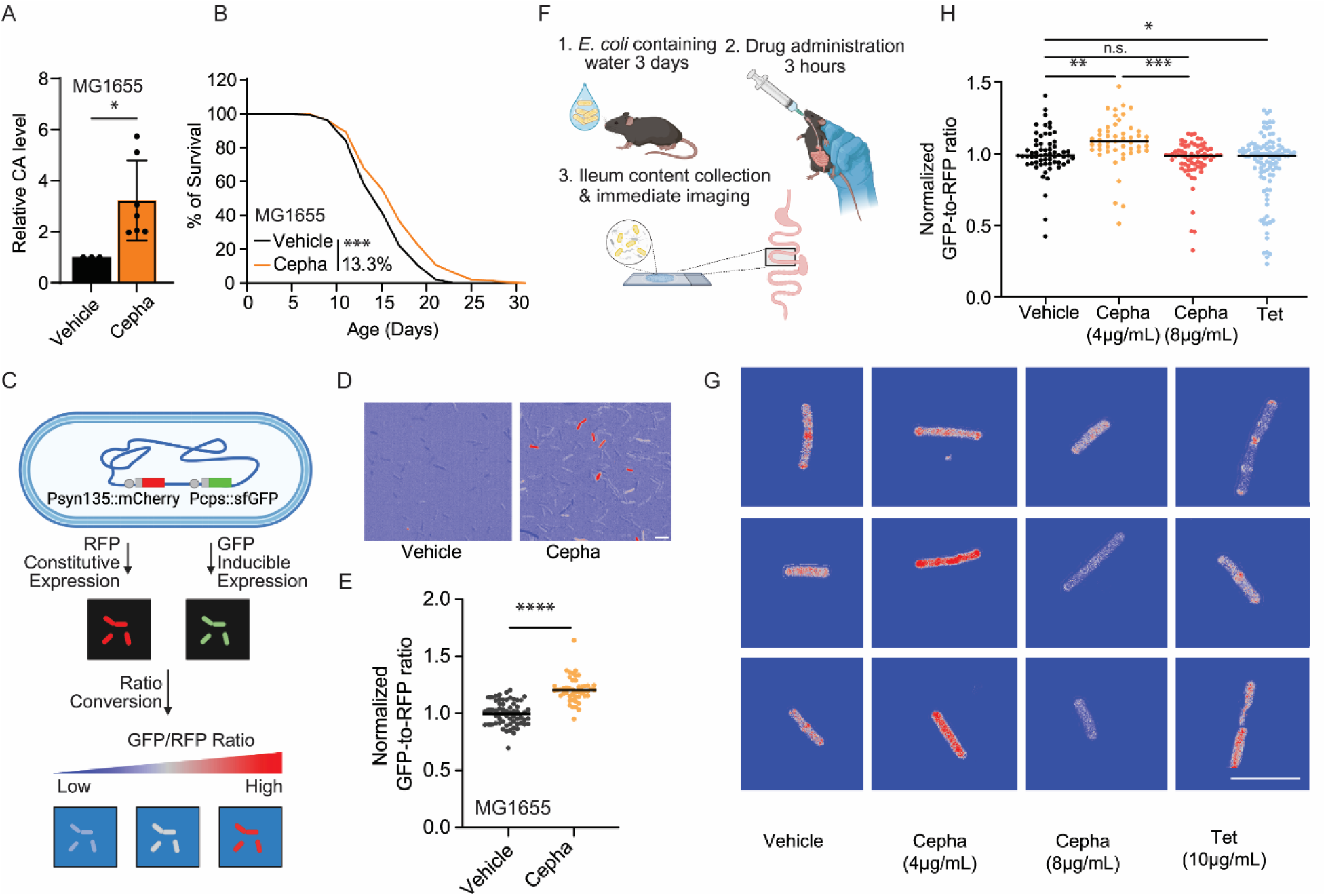
A low dose of cephaloridine up-regulates CA biosynthesis operon in the mouse gut microbiota. (A) Cepha-CAI dose-treated MG1655 *E. coli* produces increased CA levels. N>3. (B) Cepha-CAI dose-treated MG1655 *E. coli* increases *C. elegans* lifespan by 13.3%. N = 3, 60 - 100 worms per replicate. (C) Schematic diagram showing the analysis using the *cps* G-R reporter *E. coli* strain. (D) Representative ratio matric images of *E. coli* treated with vehicle or Cepha-CAI dose, scale bar=5 µm. (E) Cepha-CAI dose-treated *cps* G-R reporter strain shows a 20 % increase in GFP-to-RFP ratios compared to the vehicle-treated control. (F) Schematic diagram presenting the drug administration procedure in wildtype B6 mice. (G) Representative ratiometric images of *E. coli* from the mouse gut, scale bar=5 µm. (H) The *cps* G-R reporter *E. coli* harvested from 4 μg/mL of Cepha-treated mice show an 8 % and a 10 % increase in GFP-to-RFP ratios compared to from vehicle-treated mice and 8 μg/mL of Cepha-treated mice, respectively. The tetracycline treatment at 10 μg/mL reduced the GFP-to-RFP ratio of the *cps* G-R reporter strain by 8 %. In (A), error bars represent the mean ± standard error of the mean, * *p < 0.05* by Student’s t-test. In (B), ***p < 0.001 by log-rank test, also seen in Supplementary Table 1. In (E) and (H), black bars show the median of the data set, \**p < 0.05*, \*\**p < 0.01, ***p < 0.001, ****p < 0.0001, ns p>0.05* by Student’s t-test.

To monitor the induction of CA biosynthesis within the mouse gut, we genetically engineered the MG1655 strain to express GFP under the control of the *cps* promoter and RFP under a constitutive promoter *syn135*^22^ (*cps*G-R reporter, **Fig 2C**). In this strain, constitutively expressed RFP indicates the baseline transcription, while inducible GFP reflects the transcriptional level of the *cps* operon. We found that the Cepha-CAI dose resulted in a 20% induction of the GFP-to-RFP intensity ratio in the *cps*G-R reporter strain at 37°C (**Fig 2D, 2E**). Next, we introduced this *cps*G-R reporter strain to wild-type C57B6 (B6) mice that were raised in a specific pathogen-free environment through drinking water to examine chemical induction in the murine gut. After supplementing *cps*G-R reporter *E. coli* through drinking water for three days, we administered cephaloridine or other drugs to mice in a single low dose through oral gavage, followed by supplementing drug-containing water for 3 hours, accounting and allowing for sufficient intestinal transit time^23^ and *E. coli* turnover time^24^ (**Fig 2F**).

Leveraging the minimal intestinal absorption and utilizing low doses of cephaloridine (4 μg/mL or 9.6 μM and 8 μg/mL or 19.2 μM), we examined its effects on gut bacteria in inducing the *cps* operon. By imaging the RFP-labeled *E. coli* harvested from the luminal content of mouse ileum, we observed an 8% increase in the GFP-to-RFP ratio in mice treated with 4 μg/mL cephaloridine (**Fig 2G** and **2H**). This increase was not observed in mice treated with 8 μg/mL cephaloridine (**Fig 2G** and **2H**). In the supplement group with tetracycline (10 μg/mL) that showed no CA induction (**Fig 1A**), we observed a decrease in the GFP-to-RFP ratio instead of an increase (**Fig 2G** and **2H**). These findings suggest that a low dose of cephaloridine effectively stimulates the *cps* operon of *E. coli* in the mammalian gut microbiota.

### Chemical induction of colanic acid through new mechanisms

Next, we investigated the molecular mechanism underlying the CA induction by Cepha-CAI dose. At non-restricted temperatures below 30°C, the induction of CA biosynthesis is known to be mediated by the RCS two-component system^26^. This system involves the RcsC sensor histidine kinase situated in the inner membrane, which autophosphorylates in response to environmental cues. The RcsD phosphotransferase interacts with RcsC to transmit the phosphoryl group to the cytosolic response regulator, RcsB (**Fig 3A**). RcsA, interacting with RcsB, enhances the heterodimers’ binding with the promoter and functions as transcription factors together to up-regulate the *cps* operon, thereby stimulating transcriptional activation (**Fig 3A**).

**Fig 3.**
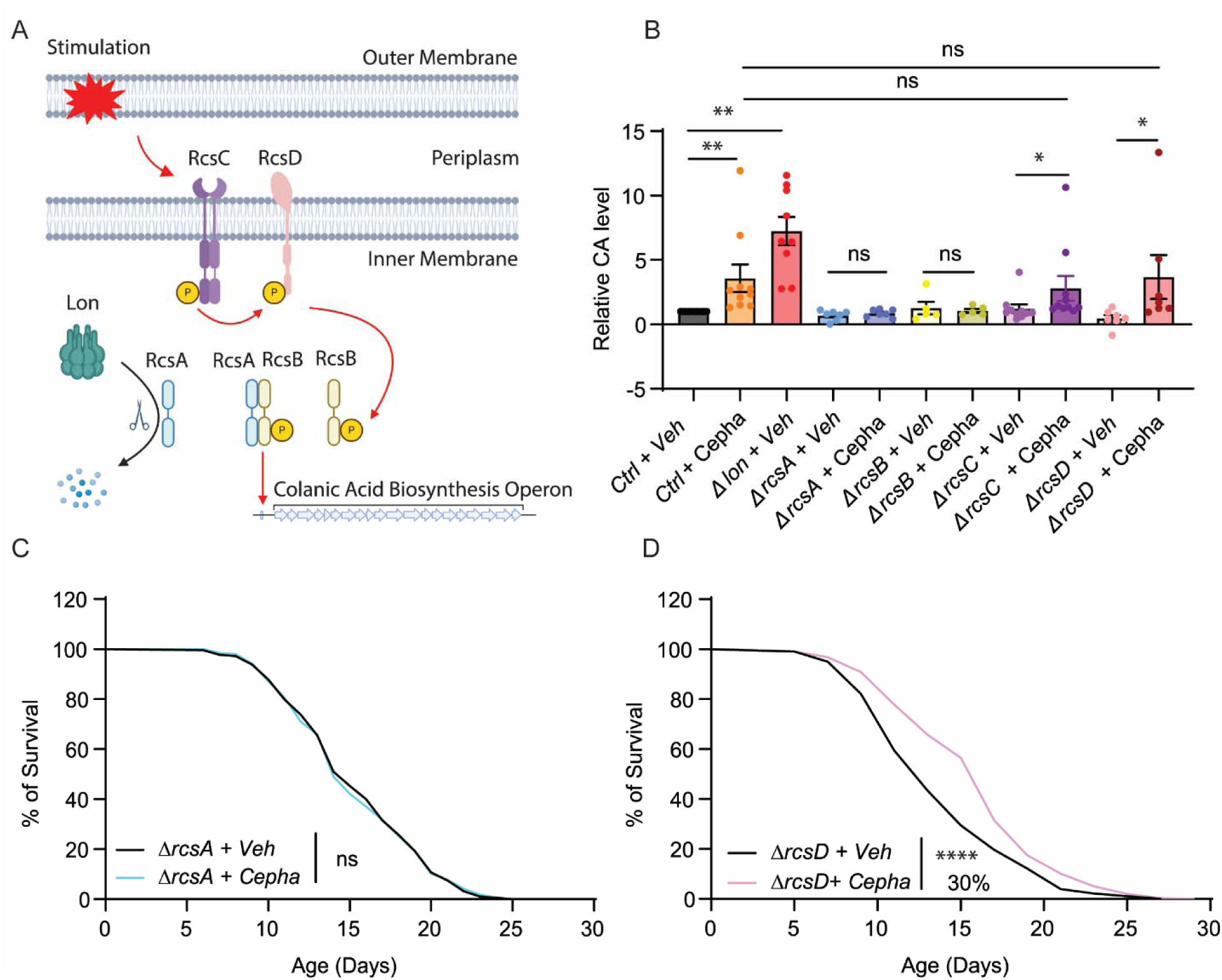
Chemical induction of bacterial CA acts independently of RCS inner membrane sensors. (A) A schematic diagram of the canonical RCS activation to induce the *cps* operon and CA biosynthesis. (B) Treatments with Cepha-CAI dose increase CA levels in the WT, Δ*rcsC*, and Δ*rcsD* mutant *E. coli* but not in the Δ*rcsA* or Δ*rcsB* mutant *E. coli*. N>3 (C) The Cepha-CAI dose-treated-Δ*rcsA* mutant *E. coli* fails to increase *C. elegans* lifespan. N = 3, 60-100 worms per replicate. (D) The Cepha-CAI dose-treated-Δ*rcsD* mutant *E. coli* increases *C. elegans* lifespan by 30%. N = 3, 60-100 worms per replicate. In (B), error bars represent mean ± s.e.m., *\*\*P < 0.01, *P < 0.05, ns P >0.05*, by two-tailed Student’s t-test. In (C) and (D), **** *P < 0.0001, ns P >0.05* by log-rank test, also seen in Supplementary Table 1.

To assess the necessity of the RCS system in the chemical induction of CA at 37°C, we treated the *E. coli* deletion mutants of *rcsA*, *rcsB*, *rcsC,* and *rcsD* with the Cepha-CAI dose. By measuring CA levels in their respective culture mediums, we found that deleting either *rcsA* or *rcsB* completely abolished the CA induction by the Cepha-CAI dose (**Fig 3B**). To our surprise, deleting either *rcsC* or *rcsD* did not suppress the induction (**Fig 3B**). These results suggest that cephaloridine relies on the RcsA-RcsB transcriptional control of the *cps* operon but operates independently of the RcsC-RcsD two-component system on the bacterial inner membrane.

Further investigation of the host lifespan in *C. elegans* revealed that, unlike cephaloridine-treated wild-type *E. coli* (**Fig 1F**), the cephaloridine-treated *rcsA* deletion mutant failed to extend the host’s lifespan (**Fig 3C**). In contrast, the cephaloridine-treated *rcsD* deletion mutant remained effective in causing lifespan extension (**Fig 3D**). These results further support that the chemical induction of CA in bacteria underlies the pro-longevity effect of low-dose cephaloridine on the host and suggest that the RcsC-RcsD inner membrane kinase complex is dispensable for the cephaloridine-mediated induction of pro-longevity CA.

### PBP1a and ZraS mediate the chemical induction of CA

We then hypothesized that the Cepha-CAI dose acts through other bacterial inner membrane proteins. Considering β-lactam antibiotics show preferential affinities to various penicillin-binding proteins (PBPs)^27,28^, we initially explored the role of these inner membrane proteins in regulating the chemical induction of CA. In *E. coli*, cephaloridine, cefazolin, and cephalothin primarily bind to PBP1a, a protein encoded by *mrcA*^29^. This contrasts with penicillin and ampicillin, which exhibit a higher affinity for PBP4, a protein encoded by *DacB*^28^ (**Fig 4A**). Notably, the Cepha-CAI dose failed to induce CA biosynthesis in the *mrcA* deletion mutant (ΔPBP1a), but this dose remained effective in the *dacB* deletion mutant (ΔPBP4) (**Fig 4B**). Thus, the low-dose cephaloridine specifically acts through PBP1a to induce CA. These findings also support that the inducing effect of the Cepha-CAI dose on CA biosynthesis is distinct from the general β-lactam antibiotic effect mediated through any PBPs.

**Fig 4.**
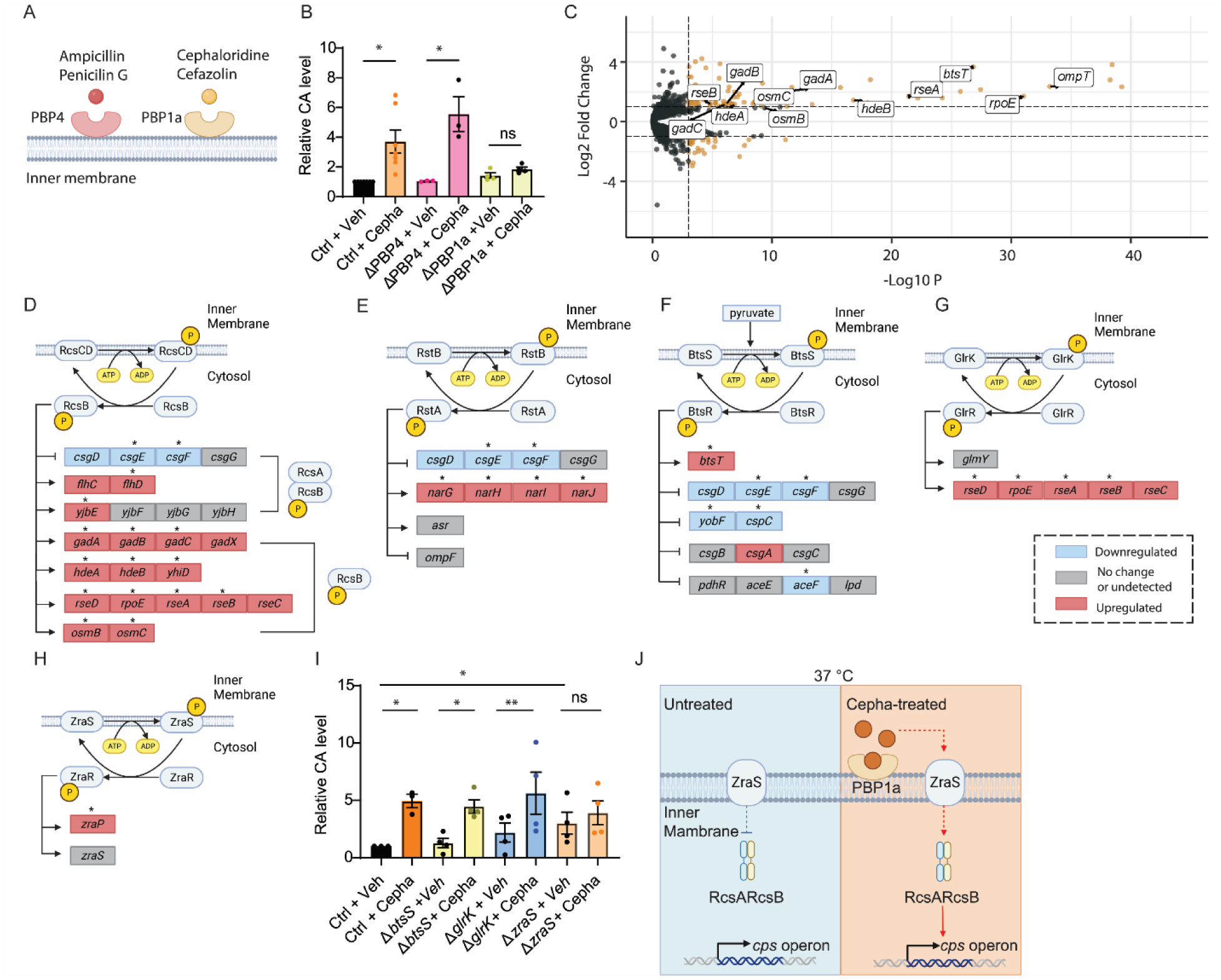
PBP1a and histidine kinase ZraS mediate CA chemical induction. (A) A schematic diagram of PBP protein binding preference of different antibiotics. (B) The loss of PBP1a encoded by *mrcA* in *E. coli* suppresses CA induction upon Cepha-CAI dose treatment. N>3. (C) A volcano plot summarizing differentially expressed genes between Cepha-CAI dose and vehicle-treated *E. coli.* (Fold change > 2, FDR adjusted P value < 0.05) (D-H) Known target genes regulated by the RcsB (**D**), BtsSR (**E**), RstBA (**F**), GlrKR (**G**), and ZraSR (**H**) two-component systems are enriched among the genes differentially expressed in the Cepha-CAI dose-treated group. Differentially expressed genes with log_2_ fold change > 0.5 in red and with log_2_ fold change < - 0.5 in blue, * *p* < 0.05; genes with - 0.5 < log_2_ fold change < 0.5 or undetectable in grey. (I) The loss of *zraS*, but not *btsS* or *glrK,* in *E. coli* suppresses the CA induction by Cepha-CAI dose. N>3. (J) Illustration of the proposed model. At 37°C, ZraS inhibits RcsAB-mediated transcription of *cps* operon. Upon cephaloridine interacting with PBP1a, it acts through the activated ZraS to up-regulate RcsAB-mediated *cps* transcription. In (B) and (J), error bars represent mean ± s.e.m., *\*\*P<0.01, *P < 0.05, ns P > 0.05* by two-tailed Student’s t-test. In (C), DESeq2 |log_2_ fold change| ≥ 0.5; P < 0.05 by two-sided Wald test (vehicle-treated control versus cephaloridine-treated control).

Next, we conducted comprehensive transcriptional profiling using wild-type MG1655 *E. coli* in response to the Cepha-CAI dose. Using RNA-seq analysis, we identified genes with significant differential expression between the Cepha-CAI dose-treated *E. coli* and the untreated control (FDR adjusted p-value< 0.05, **Fig 4C**). Among these genes, we observed an enrichment of RcsB target genes (**Fig 4D**), confirming the pivotal role of the RcsB transcription factor in the cephaloridine-mediated induction of CA. Furthermore, integrating information from transcription factor and two-component system (TCS) databases from RegulonDB led to the identification of four additional two-component systems – BtsS-BtsR (**Fig 4E**), RstB-RstA (**Fig 4F**), GlrK-GlrR (**Fig 4G**), and ZraS-ZraR (**Fig 4H**) – whose target genes exhibited enrichments among the differentially expressed candidate genes in response to Cepha-CAI dose.

To investigate the potential involvement of these TCSs, we treated the following single gene deletion mutant *E. coli* of *btsS*, *glrK*, and *zraS* with the Cepha-CAI dose at 37°C, and then measured the CA levels in their respective culture mediums. Due to its inhibited growth in the M9 culture medium, the *rstB* deletion mutant could not be evaluated in this analysis. We found that in the *zraS* deletion mutant, the Cepha-CAI dose failed to induce CA, while the inducing effect was not affected by the *btsS* or *glrK* mutant (**Fig 4I**). Notably, the *zraS* deletion mutant showed CA induction even without cephaloridine treatment, as compared to the wild-type *E. coli* (**Fig 4J**). These results suggest ZraS as the inner membrane kinase mediating the chemical-inducing effect, which typically inhibits CA biosynthesis at 37°C. Low-dose cephaloridine appears to release this inhibition, thereby triggering CA induction.

Moreover, we have systemically examined the expression levels of genes from the β-lactam resistance pathway reported by KEGG pathway analysis. We found that only *ompC,* which encodes a general outer membrane porin mediating the non-specific diffusion of small solutes, and *oppA,* which encodes the periplasmic binding protein mediating the transport of oligopeptides, are differentially up-regulated and down-regulated, respectively, in response to the Cepha-CAI dose (**Supplementary Fig 2**). These results further ensure that the Cepha-CAI dose is unlikely to stimulate the general β-lactam antibiotic stress response.

Together, we propose a model (**Fig 4J**) that ZraS inhibits RcsAB at 37°C, leading to the silencing of the *cps* operon at this temperature. Upon binding of cephaloridine with PBP1a, the activation of ZraS, which is likely associated with its autophosphorylation, triggers the transcriptional up-regulation of the *cps* operon through RcsAB.

## Discussion

In this study, we employed a xenobiotic chemical to directly target and activate a metabolic pathway of commensal *E. coli* that positively impacts the host’s fitness. We showed that a low dose of cephaloridine induces CA production from commensal *E. coli*, resulting in lifespan extension in the host *C. elegans*. This chemical induction overcomes the temperature restriction on the *cps* gene expression in the mouse intestine. Furthermore, we discovered ZraS, a histidine kinase, as a new regulator of this chemical induction in bacteria.

Interestingly, a low dose of cephaloridine induces the *cps* operon in the mouse intestine rather than a high dose, which failed to exert the same induction effect. This finding indicates that this chemical-bacteria interaction is distinct from the antimicrobial property of cephaloridine and that it is different from the drug-modifying effects of the microbiota^10,30^. Furthermore, not all antibiotics, even other β-lactam antibiotics, exert the same effect as cephaloridine in inducing the *cps* operon. This highlights the significance of the chemical specificity of cephaloridine in this regulation. Cephaloridine exhibits extremely low oral bioavailability. Its near-zero absorption in the gut has led to its discontinuation for treating infectious diseases^25^. Hence, orally supplemented low-dose cephaloridine is neither metabolized by bacteria nor directly targets eukaryotic cells. Instead, it acts as a chemical inducer, signaling through the bacterial membrane protein PBP1a and subsequently ZraS-RcsA/B to activate the *cps* operon. Based on the genetic analysis, we propose that ZraS inhibits RcsA/B activities in wild-type *E. coli*. Low-dose cephaloridine induces the *cps* operon by alleviating this inhibition. Considering the kinase activity of ZraS, it would be interesting to examine whether and how RcsAB phosphorylation is affected by ZraS in future studies.

Our findings underscore the intricate interplay between xenobiotic chemicals, bacterial metabolism, and host fitness, urging a reevaluation of conventional drug discovery paradigms primarily focusing on eukaryotic targets. Microbiota’s vast genetic diversity and metabolic pathways hold immense opportunities for developing bacteria-targeting chemical inducers or inhibitors. Further advances must rely on a systematic understanding of how microbiota-specific metabolic pathways/products influence host physiology, which will guide future drug screens focusing on microbial targets. Integrating these strategies with established microbiome-based therapeutics, as evidenced by recent studies^11^, promises a synergistic approach to tackling multifaceted health challenges.

**Supplementary Fig 1.**
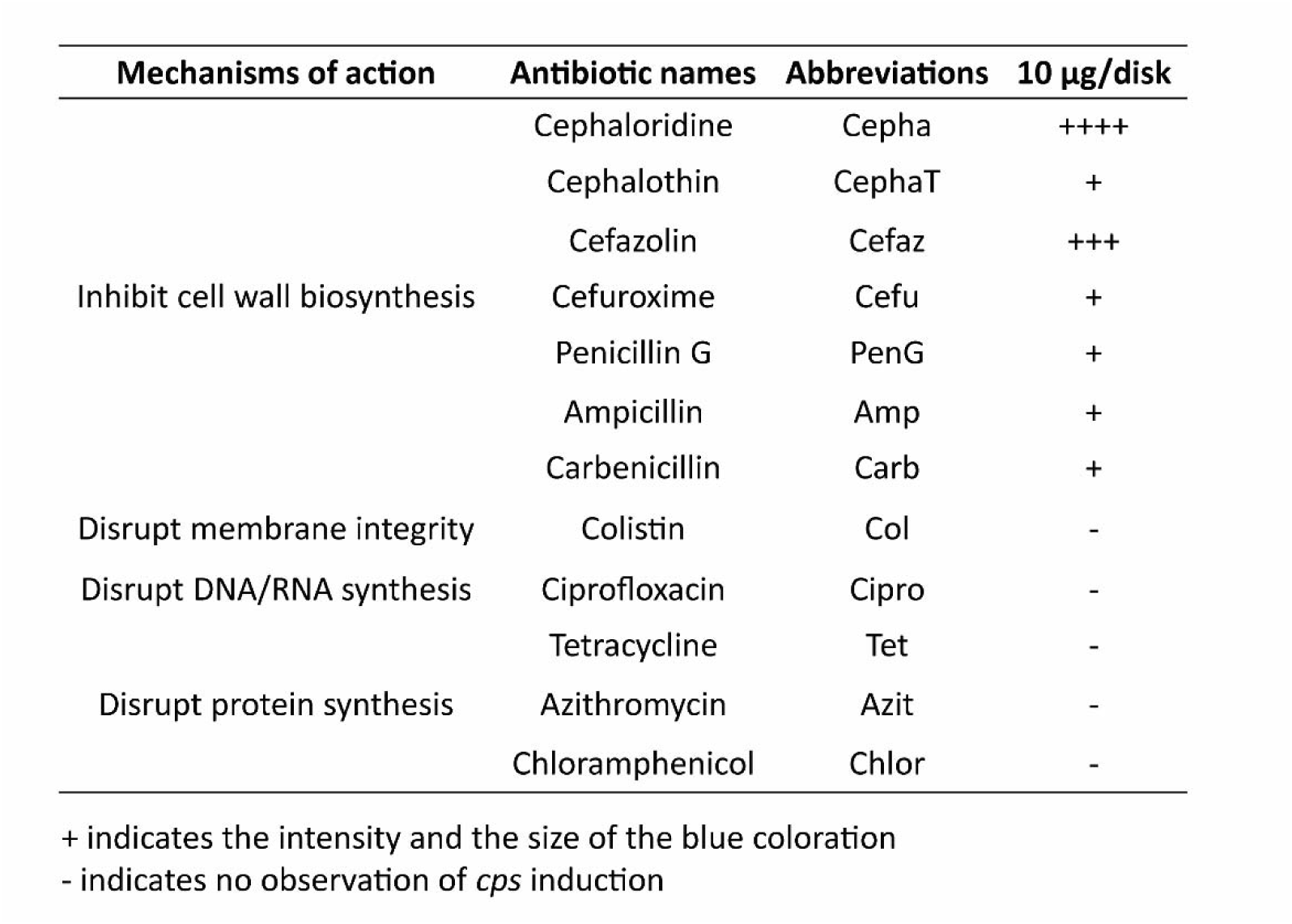
A summary of *cps* induction indicated by the intensity and the size of the blue coloration in WT BW25113 *E. coli* treated with different antibiotics.

**Supplementary Fig 2.**
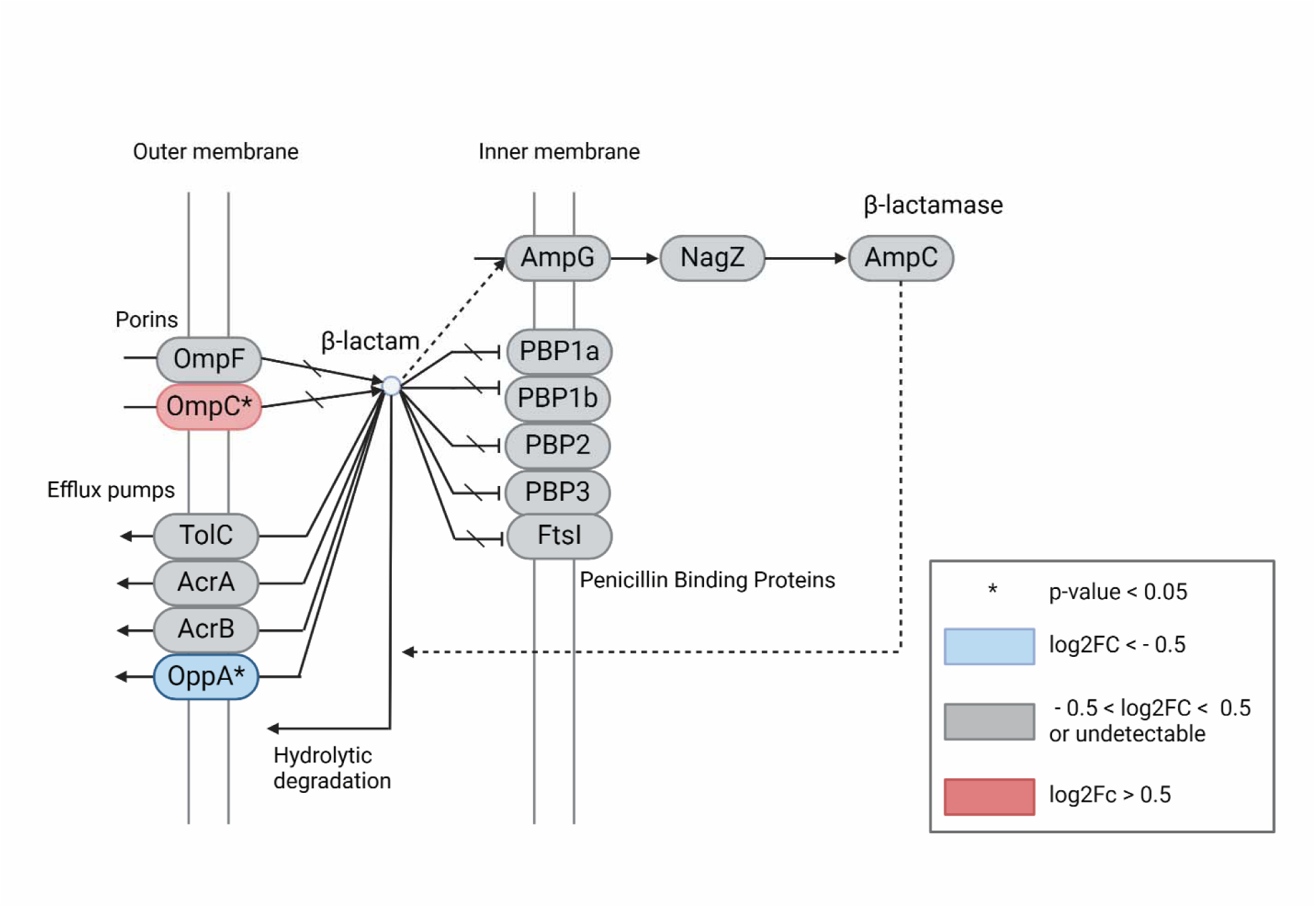
β-lactam antibiotic resistance pathway. An illustration of the β-lactam antibiotic resistance pathway in MG1655 *E. coli* adopted from KEGG (https://www.genome.jp/pathway/ko01501), which is color-coded based on the RNA-seq analysis of gene expression between Cepha-CAI dose-treated and vehicle-treated *E. coli*. Differentially expressed genes with log_2_ fold change > 0.5 in red and with log_2_ fold change < - 0.5 in blue, * *p* < 0.05; genes with - 0.5 < log_2_ fold change < 0.5 or undetectable in grey.

## Materials and Methods

### Animal and Bacteria Strains

#### 1. Bacteria strains

For <u>antibiotic screening</u> targeting *cps* transcriptional expression, we utilized the SG20781 strain (MC4100 lon+cpsB10::lacZ Mu-immλ) originally generated by S. Gottesman. The bacteria were cultured overnight at 37°C in Luria-Broth (LB) medium and evenly plated on X-gal (40 μg/mL) supplemented LB agar plates, incubated overnight.

For <u>colanic acid quantification</u>, *E. coli* Keio mutants and its parental strain BW25113 were cultured overnight at 37°C in M9 minimum medium^31^. Bacteria were then removed by centrifugation and filtration, and the culture medium was kept for colanic acid measurement.

For <u>longitudinal assays</u>, *E. coli* Keio mutants were cultivated in M9 minimum medium at 37°C for 16 hours. Subsequently, 10^7 bacteria cells from stationary phases were reinoculated in every 50mL fresh M9 medium with a low dose of cephaloridine at 37°C for another 16 hours as seeding bacterial culture. Fresh seeding bacterial cultures were made weekly. A 200 μL aliquot of the seeding bacteria culture was plated onto each 6-cm NGM plate and maintained at 20°C to transfer live worms every other day.

For <u>high-resolution imaging</u>, rph1 ilvG rfb-50 attL(Psyn135::mcherry) attHKPR218 (cps G-R reporter strain), a genetically modified strain derived from MG1655 (genotype rph1 ilvG rfb-50), was generated by Dr. Patricia Rohs. pPR218, integrated at the HK site, encodes Pcps::sfGFP and chloramphenicol resistance.

#### 2. *C. elegans* strain

*Caenorhabditis elegans* var. Bristol N2 strain was obtained from the Caenorhabditis Genome Center. The strain was maintained on standard nematode growth medium (NGM) agar plates seeded with corresponding bacteria at 20 °C. Age synchronization of *C. elegans* was achieved by isolating eggs as previously described^32^.

#### 3. Mouse colony

The female C57BL/6J mice in this study were kept on a 12-hour light/dark cycle and had access to food and water ad libitum. Animal care and experimental procedures were approved by Baylor College of Medicine’s Institutional Animal Care and Use Committee in accordance with all guidelines set forth by the U.S. National Institutes of Health.

### Disk-diffusion assay

*E. coli* strain SG20781 was used to assess antibiotic susceptibility conducted via disk diffusion previously described by the Sailer group^18^. In brief, a 16-hour overnight culture at 37°C of bacteria in LB medium was spread onto LB agar containing 0.1 mg/ml−1 5-bromo-4-chloro-3- indolyl-β-D-galactoside (X-gal). Filter papers were saturated by 100 µL of the following antibiotic solutions (cephaloridine C258600 Toronto Research Chemicals; Cephalothin, Sigma; Cefazolin, Sigma; Cefuroxime, Sigma; penicillin, Sigma; Ampicillin, Sigma; Carbenicillin, Sigma; Colistin, Sigma; Ciprofloxacin, Sigma; Azithromycin, Sigma; Tetracycline, Sigma; Chloramphenicol, Sigma) at concentrations of 10 µg mL^−1^ and air-dried. The soaked filter papers were placed onto the bacteria-seeding plates and incubated for 24 hours at 37°C.

### Colanic acid measurement

The desired bacterial strain was first inoculated in M9 minimum medium overnight at 37 ◦C as the seeding culture. The OD 600 was measured, and 10^7^ cells were reinoculated in 50mL M9 minimum medium or antibiotic-containing M9 medium for another 16 hours. After the incubation, bacterial cultures were spun down at 4,000 g at 4 ◦C for 30 min. The bacterial pellet was removed, and the culture medium was filtered through a 0.2 μm syringe filter (corning). Each 25 mL of supernatant was collected into a 50 mL conical tube, and then an equal volume of ice-cold 100% ethanol (EtOH) was used to make a final 50% EtOH for precipitation. The mixture was kept at 4 ◦C overnight. Once precipitation occurred, the liquid was carefully removed after centrifugation at 4,000 g at 4 ◦C for 50 min. The pellet was washed once with cold 80% EtOH and then air-dried in a hood. The pellet from every 50 mL culture was resuspended in 500 μL distilled H2O. The solution was sonicated in a water bath for 60 minutes at 37°C, followed by centrifugation at 4000 g at 4°C for 10 minutes. A 200 μL sample was boiled with 30 μL hydrochloric acid (320331-500mL, Sigma) for 2 hours. Subsequently, 60 μL 5M NaOH and 150 μL NaHCO3/NaOH buffer pH10 were added to reach pH 6-8, calibrated by 1M HCl or NaOH solutions. The fucose quantification assay was then carried out using a K-FUCOSE kit (Megazyme).

### Lifespan assay

Lifespan assays were performed as previously described^33^. In brief, age-synchronized *C. elegans* at the L1 stage were seeded onto designated bacteria lawns. Fresh bacteria-containing plates were made every other day for live C. elegans transfer. Once reaching adulthood as Day 0, worms were transferred to new bacteria plates. Death events were scored upon each transfer. Bagged worms and vulva protruding worms were censored through the analysis. Each assay contained 80-100 animals with 30-40 animals per 6 cm plate.

### High-resolution bacteria imaging

#### Imaging system

High-resolution bacteria imaging was performed by spinning disc microscopy using a Nikon Eclipse Ti2 system and CSU-W1 SoRa confocal scanner unit under a Nikon Plan Apo λ100x/1.45 Oil lens. The excitation wavelengths were 488nm for the GFP channel and 561nm for the RFP channel. The images were acquired using dual camera settings in the NIS-Elements imaging platform. All the raw images were processed in the ImageJ software using the same customized plugin codes generated by Dr. Chen Tao.

#### In vitro imaging

The *cps*G-R reporter strain was cultured in an M9 minimal medium starting from 10^7^ cells. Bacteria culture was harvested at an optical density of 600 of 0.6. 1 μL of the culture was directly loaded onto a 1 % low-melt agarose pad made with M9 medium^34^ for imaging.

#### In vivo imaging

Co-housed female C57B6 mice received *cps*G-R reporter *E. coli* at 1 11 10^8^ CFU per 100 mL through distilled drinking water for three days. The bacteria-containing water was prepared and replaced daily for a continuous three days. On day 3, the mice received an initial 100 μL of drug dose or sterilized water administered orally. They continued receiving drug-containing water or sterilized water for 3 hours before euthanasia. Luminal content from the ileum was collected and processed based on *in vivo* protocol^35^ immediately after euthanasia. In brief, the ileum sections were collected and then finely minced by a pair of iris scissors in 2mL 1x PBS, vortex for 1 minute, and then filtered with a sterile cell strainer (40 μm) to remove the tissue debris. The microbiota was washed three times with 1.5 mL PBS by centrifugation (15,000 x g, 3 min) and then resuspended in 0.1 mL PBS. 1 μL of the resuspension solution was directly loaded onto a 1 % low-melt agarose pad made with M9 medium for imaging.

### Transcriptional analysis

Bacterial cells were grown to a mid-exponential phase starting from an optical density of 600 of 0.03 in M9 minimal media and grown to an optical density of 600 of 0.6. 1 mL of each culture was pelleted, and RNA was extracted using RNAsnap^36^ and column purified on Zymo Clean and Concentrator columns with an off-column DNase I digestion step. Total mRNA was measured using Qubit, and the quality of RNA samples was assessed (RINe for all samples ranged from 7.9 – 8.2). Ribosomal depletion was performed with the Ribominus^TM^ Transcriptome Isolation Kit. The RNA libraries were prepared for Illumina sequencing using the Illumina stranded total RNA prep kit (Catalog No. 20040525). Libraries were sequenced on an Illumina NextSeq 550 platform with 2 x 37 cycle paired end reads. Bcl2fastq2 v2.20.0.422 was used to generate raw.fastq files, which were then filtered using FastP v0.12.4-g --poly_g_min_len 11 -l 25. TPM calculations were done on filtered.fastq’s using Salmon v1.10.0 --seqBias --gcBias --allowDoveTails to the *E. coli* MG1655 genome (NC_000913.3) with an index Kmer size of 17. DEG analysis was done on count tables using Deseq2 v1.36.0, with size factors being estimated using estimateSizeFactors() and significance assessed with nbinomWaldTest(). DEGs were visualized with pheatmap v1.0.12 and EnhancedVolcano v1.14.0 for heatmaps and volcano plots, respectively.

### Statistical analysis

For all figure legends, asterisks indicate statistical significance as follows: NS, not significant (P > 0.05), *P < 0.05, **P < 0.01, ***P < 0.001, and ****P < 0.0001. Data were obtained by performing independently at least three biological replicates unless specified in the figure legends. No statistical method was used to pre-determine the sample size. No data were excluded from the analyses. Two-tailed Student’s t-test or one-way or two-way analysis of variance (ANOVA) with Holm–Sidak corrections were used as indicated in the corresponding figure legends. N indicates the number of biological replicates. n indicates the number of animals or technical replicates within each biological replicate. For survival analysis, statistical analyses were performed with SPSS software (IBM) using Kaplan–Meier survival analysis and log-rank test, as well as using GraphPad Prism 10 survival analysis. For RNA-seq, a two-sided Wald test in R package DEseq2 was used. Figures and graphs were constructed using BioRender.com, ImageJ, GraphPad Prism 10 (GraphPad Software), and Illustrator (CC 2019; Adobe).

## Data Availability

Sequencing data is available through the Sequence Read Archive under the accession number PRJNA1111054.

All relevant data generated or analyzed in this study are included in this manuscript and/or its supplementary information as source data or can be obtained from the corresponding authors upon request.

## Code Availability

The code utilized for high-resolution bacteria imaging is available on GitHub (https://github.com/huguo0519/Cepha-project.git)

## Acknowledgments

We thank Dr. J. Petrosino, Dr. T. Palzkill, Dr. W. Craigen, and Dr. D. Moore for invaluable guidance and discussions on this project. We express strong gratitude to Dr. P. Roh for generating the *cps*G-R reporter strain and Dr. T. Chen for generating the analytical codes for bacteria fluorescence intensity analysis. The research of this work is supported by Howard Hughes Medical Institute (M.C.W.) and is funded by NIH DP1 AI152073 to C.H.

## Author contributions

G.H. and M.C.W. conceptualized the project, designed the experiments, and contributed to data visualization. G.H performed the experiments and data analysis, except the bacterial transcriptional analysis, in which the library assembly was performed together with A.X.W., and RNAseq and data analysis were performed by M.B.C. and supervised by C.H. G.H coordinated the project as manager and prepared the original manuscript draft. G.H., M.S., A.X.W., M.B.C., C.H., J.W., reviewed and edited the manuscript. M.C.W supervised the project.

